# Providing Insights into Health Data Science Education through Artificial Intelligence

**DOI:** 10.1101/2024.03.22.586308

**Authors:** Narjes Rohani, Kobi Gal, Michael Gallagher, Areti Manataki

**Affiliations:** Usher Institute, University of Edinburgh, UK; School of Informatics, University of Edinburgh, UK; Moray House School of Education and Sport, University of Edinburgh, UK; School of Computer Science, University of St Andrews, UK

**Keywords:** Health data science, Artificial intelligence, Learning analytics, Learning strategy, Learning tactic, Learning engagement, Health informatics, Medical education, Educational data mining

## Abstract

**Background:** Health Data Science (HDS) is a novel interdisciplinary field that integrates biological, clinical, and computational sciences with the aim of analysing clinical and biological data through the utilisation of computational methods. Training healthcare specialists who are knowledgeable in both health and data sciences is highly required, important, and challenging. Therefore, it is essential to analyse students’ learning experiences through artificial intelligence techniques in order to provide both teachers and learners with insights about effective learning strategies and to improve existing HDS course designs.

**Methods:** We applied artificial intelligence methods to uncover learning tactics and strategies employed by students in an HDS massive open online course with over 3,000 students enrolled. We also used statistical tests to explore students’ engagement with different resources (such as reading materials and lecture videos) and their level of engagement with various HDS topics.

**Results:** We found that students in HDS employed four learning tactics, such as actively connecting new information to their prior knowledge, taking assessments and practising programming to evaluate their understanding, collaborating with their classmates, and repeating information to memorise. Based on the employed tactics, we also found three types of learning strategies, including low engagement (Surface learners), moderate engagement (Strategic learners), and high engagement (Deep learners), which are in line with well-known educational theories. The results indicate that successful students allocate more time to practical topics, such as projects and discussions, make connections among concepts, and employ peer learning.

**Conclusions:** We applied artificial intelligence techniques to provide new insights into HDS education. Based on the findings, we provide pedagogical suggestions not only for course designers but also for teachers and learners that have the potential to improve the learning experience of HDS students.

## 1 Background

In recent decades, data science and Artificial Intelligence (AI) techniques have shown promising applications in the field of medicine, leading to the emergence of an interdisciplinary field named Health Data Science (HDS) [1, 2]. The availability of massive amounts of clinical and patient data has provided the opportunity for utilising computational tools to aid healthcare professionals in decision-making [3]. The application of data science in medicine has the potential to discover breakthrough findings that could improve global health through the early detection of diseases and personalised treatment recommendations [1, 4–6].

Unfortunately, despite the high demand for data-literate healthcare specialists, according to the National Academy of Medicine, training students in this field can be challenging [7]. This difficulty is exacerbated by a variety of factors, such as the complexity of teaching both medical and computational concepts to students with diverse backgrounds, and the uncertainty over students’ learning preferences [2, 8, 9]. Consequently, both instructors and learners find it challenging to teach and learn HDS courses [8–11]. Conducting research to analyse HDS courses through AI techniques is thus necessary to understand how students regulate their learning, and identify areas for improvement. This information can be leveraged by instructors to facilitate better learning outcomes for HDS students and to design courses that are aligned with the HDS students’ needs and preferences [12, 13].

Only a few studies [14–16] have been conducted to explore the learning preferences of students in HDS-related courses, specifically in the fields of bioinformatics [17] and precision medicine [15]. All of these studies relied on self-reported data, which can be biased and may not accurately reflect the true behaviours of students [18, 19]. For example, Micheel *et al.* [15] conducted a survey study to discover the learning preferences of precision medicine healthcare professionals. Their findings showed that 80% of participants had multiple learning preferences. The largest group (39% of the participants) preferred a combination of watching, listening, and reading, whereas 19% of participants preferred a combination of watching and reading. The authors compared an intervention group that was exposed to a personalised course based on their learning preferences, with a control group that received standard training. The intervention group achieved significantly higher scores in both past-test and follow-up tests, suggesting that providing a customised course based on students’ preferences can significantly improve their learning outcomes. They also showed that the learning preferences of HDS students differ from those of medical students.

In another study, Holtzclaw *et al.* [16] explored four dimensions of learning preferences, including sensing/intuiting, visual/verbal, and active/reflective. The survey results indicated that the majority of bioinformatics students had visual (82%) and sequential (75%) learning preferences. The authors also conducted further analysis using pre- and post-course surveys, which confirmed their conclusions. According to this study, teaching genetic concepts through the use of visualisation techniques, such as diagrams and plots, was more effective for bioinformatics learners.

Although self-reported data can provide insight into students’ preferences and opinions, such information may be limited and biased due to several reasons [20, 21]. First, the amount of data that is collected is not large enough to support rigorous statistical analysis. Second, the information reported by students may be influenced by their self-perceptions, which may not always align with their actual behaviours. Third, self-reported data may be subject to accidental or deliberate misreporting. We posit that applying AI techniques on log data collected from students’ actual experience in a course can provide a more representative dataset for analysis [18, 22].

Research findings in various disciplines suggest that the analysis of log data using AI is a reliable approach to discovering students’ learning behaviours, such as their learning tactics and strategies [23–25]. For example, Jovanovic *et al.* [25] analysed the log data from an engineering course and discovered four learning tactics and five learning strategies by using sequence mining and Agglomerative Hierarchical Clustering (AHC). Maldonado-Mahauad *et al.* [26] also applied a process mining technique to analyse log data from three Massive Open Online Courses (MOOCs) in education, management and engineering. They discovered four learning patterns from students’ interactions with the course content and identified three groups of learners: “Comprehensive” learners who followed all course structure steps, “Targeting” students who focused on a specific set of activities that helped them pass the assessments, and “Sampling” learners with less goal-oriented strategy and low engagement.

Subsequently, Matcha *et al*. [24] analysed the log data from biology, Python programming and computer engineering courses using process mining and the Expectation Maximisation algorithm, which resulted in the discovery of various learning tactics in each course. Then, they applied AHC to the frequency of using the identified learning tactics by each student, which yielded three groups of students, namely low, moderate, and high engagement. Similarly, Crosslin *et al.* [27] used the same method to identify five learning tactics and four learning strategies employed in an online college-level history course.

Although AI methods have been employed to analyse the students’ learning experiences in a few disciplines [24–27], there is no research that analyses HDS students’ learning strategies using the power of AI algorithms. Given the fact that there is no data-driven insight into the learning behaviours of HDS students, multiple papers have emphasised the importance of conducting studies to analyse HDS education and learners’ experiences [28–31].

This paper directly addresses the above shortcoming by using AI techniques to analyse students’ interactions in an HDS MOOC with over 3,000 learners. We used statistical methods to explore students’ engagement with different types of educational resources (video lectures, reading materials, and so on) across three performance groups (students with low, moderate, and high performance). We also explored students’ engagement with different HDS topics covered in the course (e.g., medical image analysis, Python programming, and network biology) to identify topics that were most interesting or difficult for the students.

We identify a hierarchy of student activities in the course, ranging from low-level activities (e.g., watching videos, answering a question in a forum), mid-level learning tactics (e.g., collaborating with other students), and high-level strategies (e.g., deep learners).

We used statistical and AI methods to investigate the following research questions.

- RQ1 - What type of educational resources in the HDS MOOC did the students engage with?
- RQ2 - What health data science topics in the HDS MOOC did the students engage with?
- RQ3 - What learning tactics and strategies did the students employ in the HDS MOOC?
- RQ4 - Is there any association between students’ learning tactics and strategies and their performance?

With respect to RQ1, we found that overall, there were no large differences in engagement between readings and lecture videos, but students who achieved higher final grades engaged more than other students in all types of resources, especially in quizzes, labs, and projects. Regarding RQ2, among the taught topics, students were more engaged with Python programming and Sequence Processing.

With respect to RQ3 and RQ4, we identified the following four prevalent learning tactics employed by the HDS students: Elaboration – actively connecting new information to existing knowledge, Problem-solving – solving assessments and programming questions for better understanding, Peer learning - collaborating with peers to share knowledge, and Rehearsal – repeating information for better retention. Based on the frequency of using the identified learning tactics, we discovered three types of strategies employed by students, which are directly aligned with educational theory: low engagement (Surface learners), moderate engagement (Strategic learners), and high engagement (Deep learners). We found that the elaboration tactic had the highest correlation with overall student performance, and deep learners who had high final grades used this tactic more. Based on our findings, we provide pedagogical recommendations for course designers, teachers, and learners in HDS that can potentially improve HDS education.

## 2 Methods

In this section, we will provide a general description of the HDS MOOC, the student population, and the data that was available for analysis. Then, we will detail the methodology used to address each of the research questions.

### 2.1 Course and Participants

The study is based on the Data Science in Stratified Healthcare and Precision Medicine (DSM) MOOC offered by the University of Edinburgh on Coursera [32]. We focused on the period between April 2018 and April 2022, and we analysed the log data of 3,527 learners who engaged with at least one learning activity (see supplementary material for considered learning activities). The course completion rate for these students is 38%.

Demographic information of students shows that 37% were male, 28% were female, and 35% did not report their gender. Regarding their educational background, 15% held a master’s degree, 12% had a bachelor’s degree, 7% held a doctorate, and the remaining had a lower degree or did not report their level of educational attainment. There is a good location spread, with 34% of students based in America, 30% in Asia, 22% in Africa, 11% in Europe, and 2% in Oceania. This study used anonymised data and received institutional ethical approval.

DSM is an intermediate-level MOOC with a total of 43 videos, 13 reading materials, five quizzes, six discussion forums, one programming assignment and one peer-reviewed project assignment. The course covers the following five topics/weeks:

1. Course Introduction and Introduction to Programming
2. DNA Sequence Processing and Medical Image Analysis
3. Biological Network Modelling, Probabilistic Modelling, and Machine Learning
4. Natural Language Processing and Process Modelling in Medicine
5. Graph Data, and Ethical and Legal Aspects

Each topic includes case studies, which are optional interview videos with specialists discussing real-world HDS projects and their research areas. The course assessment includes a quiz for each topic, as well as a programming assignment for the third topic and a peer-reviewed project on the last topic of the course. Final student grades are calculated (out of 100) using a weighted average of all quiz and assignment scores, with each quiz worth 10%, the programming assignment worth 20%, and the peer-reviewed assignment worth 30%.

### 2.2 Data Analysis

Two-sided t-tests as well as Cohen’s D effect size were used to investigate any differences regarding engagement with different types of educational resources and course topics (RQ1 and RQ2).

For RQ3, several AI techniques were used to analyse the log data of the DSM course to uncover the learning tactics employed by the students. Following the approach by Matcha et al. [24], the click-stream data from the beginning to the end of the course was divided into different learning sessions. A learning session represents a consecutive series of learning actions performed by a student within one login into the learning platform. After pre-processing the learning sessions (this included considering two consecutive sessions with a time gap less than 30 minutes as one session), 44,505 learning sessions were identified. Process mining and clustering methods were employed to detect the learning tactics. In particular, the probability of switching between different learning actions was estimated with the use of First-Order Markov Models (FOMMs) as implemented in the pMinerR package [33]. The number of possible learning tactics (no. tactics = 4) was determined based on a hierarchical clustering dendrogram. To identify the learning tactics, the Expectation-Maximisation (EM) algorithm was applied to the calculated transition probability matrix.

Students often use a set of learning tactics while interacting with the course. Therefore, a learning strategy is defined as the goal-driven usage of a collection of learning tactics with the aim of acquiring knowledge or learning a new skill [24, 27, 34, 35]. Based on previous work [24], the frequency of using each learning tactic by each student was calculated as a measure to group students into categories of learning strategies. The learning strategies were identified through agglomerative hierarchical clustering with the Ward algorithm. The potential number of clusters (no. groups=3) was determined based on the height of the dendrogram.

To answer RQ4, the Pearson correlation and the two-sided t-test were used to check for any correlation between different learning tactics and students’ grades in different assessments. The Kruskal-Wallis test was also used to test the association between students’ learning strategies and performance. In other words, we explored both the association of a single learning tactic and the collection of learning tactics (learning strategy) with students’ final grades because it is useful to know whether any single tactic was more correlated to a higher final grade. Further information about the methodology can be found in the supplementary material.

## 3 Results

In this study, similarly to previous work [24, 27], we considered the number of clicks made by a student in the learning platform as a metric to assess their level of engagement. Also, following the course instructor’s recommendation, the students were categorised into three performance levels. **LP**: Low-Performance (final grade below 50, representing 62% of students), **MP**: Moderate-Performance (final grade between 50 and 80, representing 21% of students), and **HP**: High-Performance (final grade of 80 or higher, representing 16% of students). These performance categories were used in the next investigations to analyse the level of engagement of students in each performance group.

### 3.1 Engagement with Different Types of Educational Resources

***Table 1*** presents the relative number of visits to each type of educational resource. To provide a fair comparison between students’ engagement with each type of educational resource, we only measured the number of visits to the parent pages of each type of educational resource (on the Coursera platform there is a parent page for each resource before students go through the children/link pages of those resources).

**Table 1.** Relative engagement in each type of educational resource. The “Count” column indicates the number of materials of each type available in the course. The “Avg. no. visits” column shows the average number of clicks per student per type of resource. The “Relative no. visits” is calculated by dividing the “no. visits” by “count” for normalisation purposes.

| Resource Type | Count | Avg. no. visits | Mean Relative no. visits | Std. of relative no. visits |
| --- | --- | --- | --- | --- |
| Peer-reviewed project | 1 | 5.48 | **5.48** | 10.74 |
| Quiz | 6 | 9.77 | 1.95 | 1.83 |
| Programming lab | 7 | 8.63 | 1.23 | 1.55 |
| Video lecture | 43 | 47.31 | 1.1 | 0.95 |
| Reading | 13 | 12.72 | 0.9 | 0.65 |
| Discussion | 5 | 2.53 | 0.51 | 1.42 |

The results in ***Table 1*** show that, as expected, the relative number of visits to assessments (peer-reviewed project and quiz) is considerably higher than those to other learning materials (lectures, readings, labs, and discussions), demonstrating that students spend more time on assessments, particularly on the peer-reviewed project (Relative no. visits=5.48). This could be because the peer-reviewed project needs more effort and accounts for 30% of the final grade. Therefore, it is not surprising that students visited this resource more than others.

Additionally, based on ***Table 1***, there is a small difference (t = 3.29e-74, p = 3.29e-74, d = 0.24) in the relative number of visits to video lectures (mean =1.1) and reading materials (mean = 0.9). Similarly, the relative number of visits to programming labs is a bit higher than those to lectures and reading materials (pl = 9.89e-09, tl = 5.7466, pr = 7.64e-44, tr = 14.0826, dl =0.10, dr = 0.28). It can be inferred that students found programming labs either a bit more interesting or challenging, leading to more visits. However, the D effect size of the difference between lab and lecture is less than 0.2; therefore, the difference is not big.

It is worth noting that after measuring the average number of visits for children (linked) and parent (main) webpages related to each educational resource, students had relatively high engagement within each resource material. As an example, even though discussion forums have fewer visits, students exhibited higher engagement (mean no. visits=11.29) and movement between posts within the forums compared to their visit to parent pages of discussion forums. However, to have a fair comparison between different types of educational resources, only the number of visits to the parent pages of each resource was measured (refer to ***Table 1***).

We explored the engagement with different types of educational resources for the HP, MP, and LP groups (**Fig. 1**). For the HP group, engagement in the project (Relative no. visits=18.16) was higher (d = 0.63, t = 28.085, p = 8.56e-110) than in the quiz (Relative no. visits=3.37). Additionally, their engagement in the lab (Relative no. visits=2.14) was slightly (+1.5 clicks) higher than in the lectures, readings, and discussions. Conversely, for the LP group, the quiz (Relative no. visits=1.14) had the highest relative number of visits.

**Fig. 1.**
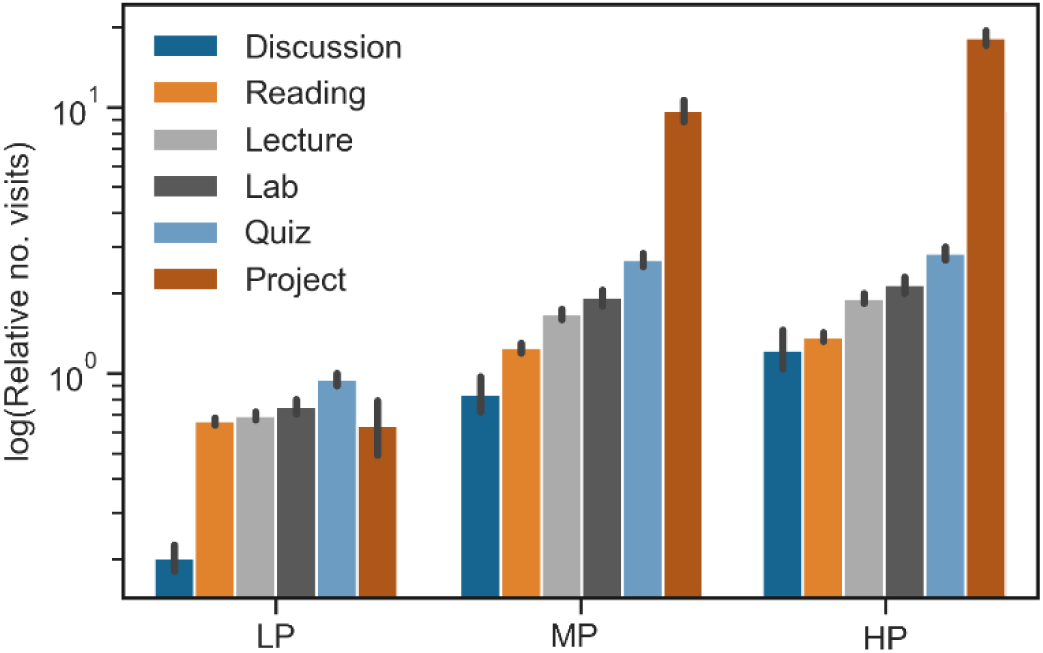
Relative no. visits of parent pages of each type of educational resource. The figure shows log of the relative number of visits for each educational resources by each group of students. LP: low performance students, MP: moderate performance students, HP:

It can be inferred that, overall, there were no large differences in engagement between readings and lectures among HDS students, but students who achieved higher final grades (both HP and MP) visited more time all types of resources, especially in project. The discussion forums were also used more by HP students than by LP students (LP’s relative no. visits = 0.20, HP’s relative no. visits = 1.22, t = 10.470, p = 2.52e-23, d = 0.28).

### 3.2 Engagement with Different HDS Topics

The analysis regarding engagement with different HDS topics was based on two measures: 1) relative average video lecture watching time, and 2) click-based interaction with videos.

**Fig. 2** shows the average watching time, standardised by dividing by the length of each video lecture, for each topic. The bar plot demonstrates that students dedicated more time to “Introduction to Programming” and “Sequence Processing” compared to the other topics. Moreover, the figure shows that the engagement of students with the initial topics were more than the final topics.

**Fig. 2.**
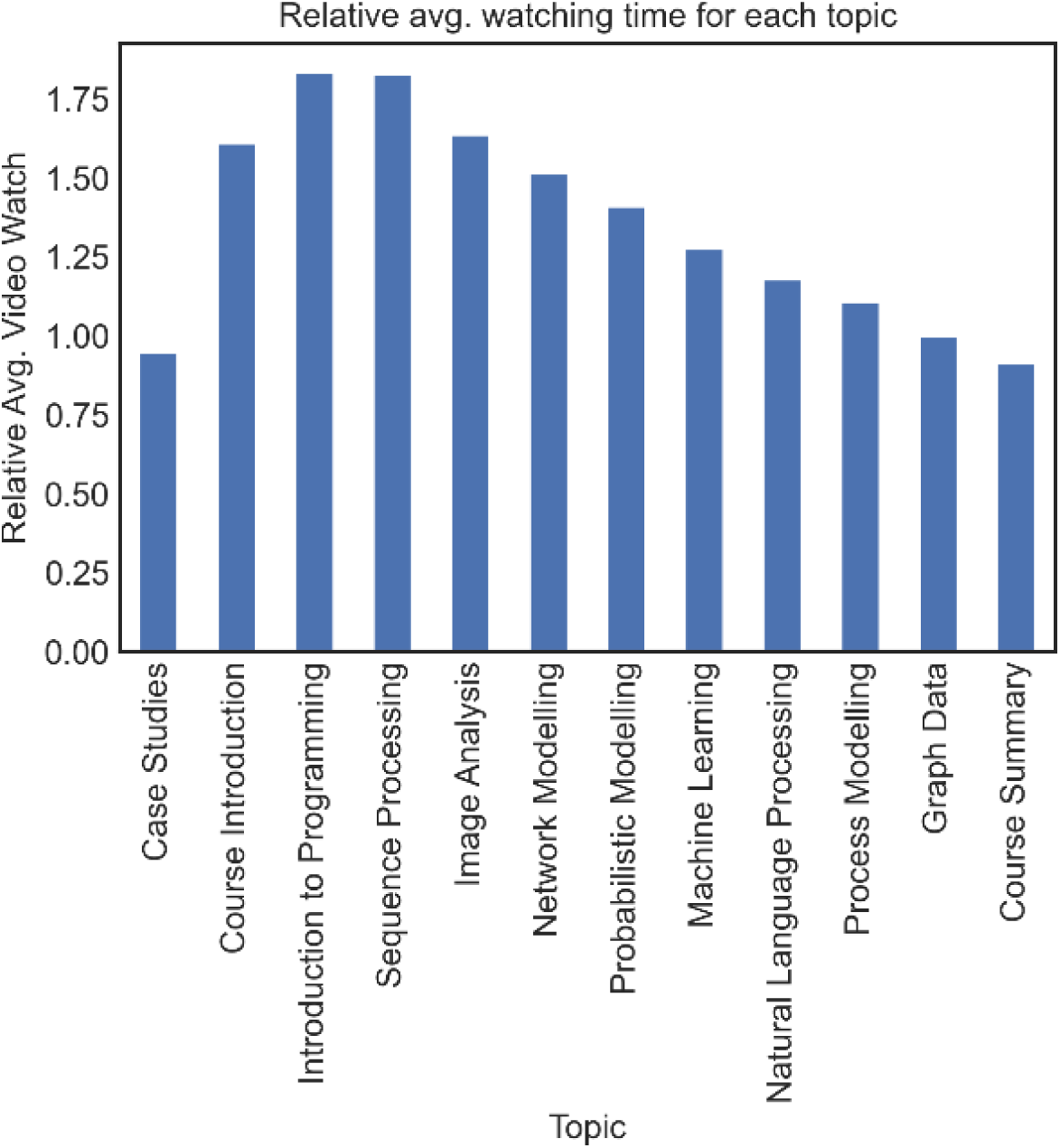
The relative average watching time of each video lecture for students. The relative video watch was calculated by dividing the average watched time by the total length of each video lecture.

Upon performance group-based analysis of video lecture watching time, we also found that HP students showed higher engagement in extra-curricular activities by spending more time watching the case studies (avg. watching time for HP = 0.62, avg. watching time for LP = 0.33). These case studies are optional interview videos in which HDS researchers and practitioners discuss their research projects. This finding is supported by previous research [24, 36, 37] that high achievers not only focus on the required syllabus but also aim to gain a deep understanding of the topics, often going beyond the syllabus.

The analysis of the students’ video interaction data reveals that students played, paused, and went forwards and backwards (seek) in the Introduction to Programming topic more than in other topics (as shown in Error! Reference source not found.). Our interpretation is that students potentially found this topic to be challenging, and therefore they rewatched certain parts to improve their understanding.

Apart from the Introduction to Programming topic, students had high click-based engagement with the Sequence Processing, Image Analysis, Network Modelling, and Machine Learning topics. It is also worth mentioning that although the case study videos were not mandatory, they achieved relatively high engagement according to the course instructor’s point of view. Based on the unexpected engagement of students with the case studies, one can infer that HDS students were interested in practical knowledge and real-world examples [24, 36, 37].

### 3.3 Learning Tactics

Based on existing literature [24, 25, 27], a learning tactic is defined as a series of actions that a student carries out to fulfil a specific task in their learning procedure. After analysis of the students’ learning sessions, we discovered that the DSM students employed four learning tactics: Elaboration; Programming and Problem-solving; Peer learning; and Rehearsal. These learning tactics are in line with the Motivated Strategies for Learning Questionnaire (MSLQ) learning theory, and we named the data-driven tactics based on this learning theory [38].

**Elaboration** is the longest (median 25 actions per session) and the most frequently used (45% of all learning sessions) learning tactic by DSM students (see ***Table 2***). The dominant learning actions in this tactic are “video play” and “pause” (**Fig. 4.a**). We can infer that this tactic primarily focuses on learning theoretical concepts, rather than programming practice and assessment participation. Students might have paused video lectures to reflect on their acquired information by taking notes, thinking about the new knowledge, or connecting concepts to their prior knowledge. This is in line with the MSLQ theory, which explains the elaboration tactic as a cognitive process in which students actively reflect on and connect new information to their prior knowledge, which can enhance their learning outcomes [38, 39]. Our results also confirm that this tactic has a positive correlation with student performance. The positive Pearson correlation between the number of times students used the elaboration tactic and their final performance (r = 0.38, p = 1.43e-113) provides evidence of the effectiveness of this learning tactic.

**Table 2.** Discovered learning tactics for HDS students.

| Tactic | Percentage | Length | Dominated learning actions | Description | Association with Final Performance |
| --- | --- | --- | --- | --- | --- |
| Elaboration | 45% of all learning sessions | Median 25 actions per session | Video pause and replay | This tactic includes using various learning actions; however, the dominant learning actions are video replay and video pause. | $r=0.38$, $p=1.43e-113$ |
| Programming and problem-solving | 24% of all learning sessions | Median 4 actions per session | Lab and quiz | As dominant learning actions in this tactic are lab and quiz. | $r=0.35$, $p=1.30e-92$ |
| Peer learning | 19% of all learning sessions | Median 5 actions per session | Peer-reviewed and discussion | The peer-reviewed project and discussion are the most used learning actions in this tactic. | $r=0.55$, $p=3.04e-260$ |
| Rehearsal | 12% of all learning sessions | Median 9 actions per session | Video seek and revisit | The most frequent learning actions in this tactic are video seek and video revisit. | $r=0.19$, $p=8.82e-29$ |

**Fig. 3.**
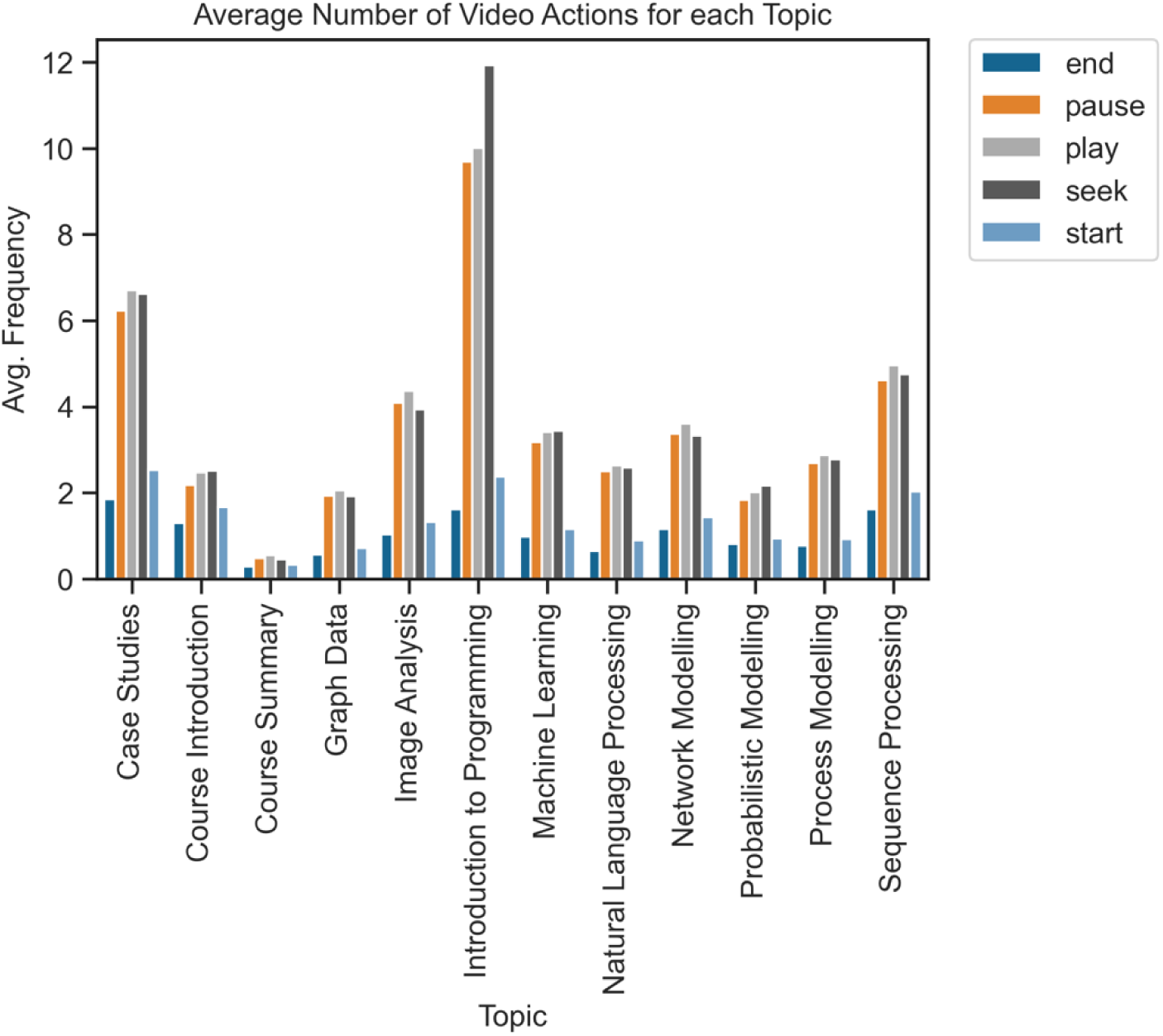
Average frequency of using each video action per student during watching each topic. “end” indicates watching a video until the end. “pause” means the student paused the video. “seek” means going forwards or backwards in a video. “play” indicates replaying a video after a pause. “start” means starting a video from the beginning. The topics in the x-axis are listed based on the order of teaching in the course (with the exception of Case Studies, which are spread across the course duration).

**Fig. 4.**
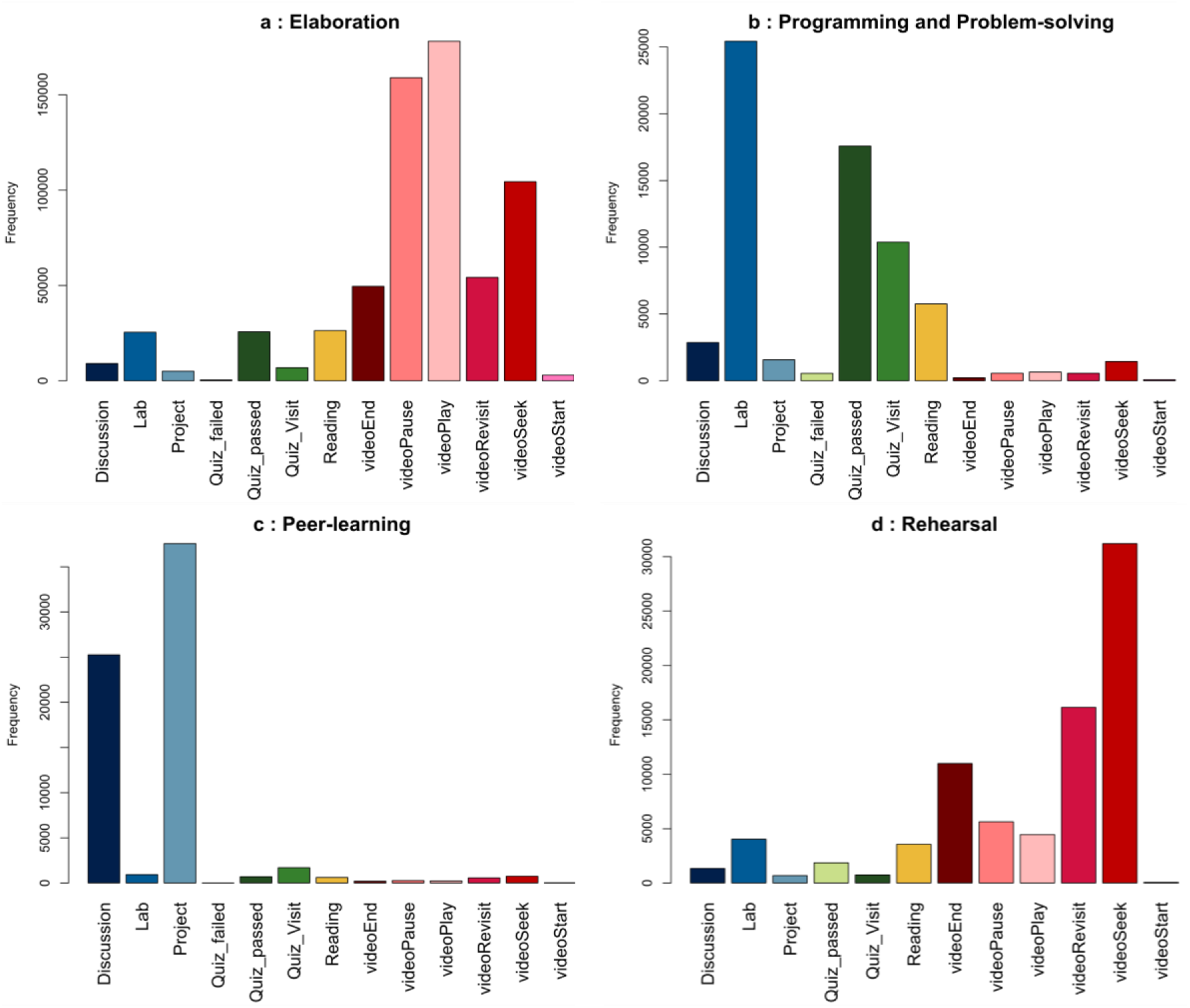
Frequency plot of each learning tactic, showing how many times each learning action was used in that tactic.

**Programming and Problem-solving** is the second most frequently used learning tactic (24% of all sessions). The dominant learning actions in this tactic are related to labs and quizzes (**Fig. 4.b**). It can be concluded that DSM students employed this tactic for applying their acquired knowledge in solving programming labs as well as the programming assignment. It seems that they participated in quizzes to assess their knowledge and solve problems.

The positive Pearson correlation (r = 0.35, p = 1.30e-92) between the frequency of using this tactic and their final performance suggests that this tactic is effective for achieving a high final grade. In conclusion, the students who practice programming more often tend to be more successful, which is consistent with [14]. This is because programming skills are an essential aspect of health data science and programming practice helps students develop critical thinking and problem-solving, which are essential skills for analysing health data. Therefore, students who invest time in practicing programming tend to have a better understanding of the material and perform better in the course assessments.

**Peer learning** is the third most frequently used learning tactic (19% of all sessions), in which DSM students engaged with discussion forums to ask questions, read others’ discussions, reply to peers, and solve the peer-reviewed project. The dominant learning actions in this tactic are the peer-reviewed project and discussion (**Fig. 4.c**). The correlation analysis also shows a positive correlation between the number of times students used peer learning and their final performance (r = 0.55, p = 3.04e-260), which is stronger than for the other learning tactics. This is not surprising, as 30% of the final grade is related to the peer-reviewed project, and students who engage more in peer learning are expected to achieve a higher grade. However, the correlation analysis between the number of times students used the peer learning tactic and their average grade in quizzes also shows a positive correlation (r = 0.46, p = 5.29e-172). This supports the conclusion that the peer learning tactic is one of the most effective learning tactics, and this is in line with existing research that has shown that peer learning can lead to enhanced motivation, increased engagement, and improved learning outcomes [14, 38, 39].

**Rehearsal** is the least used learning tactic by DSM students (12% of sessions), and it is focused on acquiring theoretical knowledge. Although both Rehearsal and Elaboration learning tactics involve learning theoretical knowledge and mostly watching video lectures, the dominant learning actions in Rehearsal are video seek and video revisit actions (**Fig. 4.d**). This suggests that students may have employed this learning tactic by reviewing certain parts of the video lectures instead of deeply understanding concepts through reflection and note-taking. In other words, Rehearsal is a simple tactic for memorising and superficially looking at learning materials. Based on prior research, the Rehearsal tactic may result in temporary retention of information rather than long-term retention [38]. As a result, some studies have indicated that the impact of this tactic may be restricted to low-level learning outcomes [39]. Although our Pearson correlation analysis shows a positive correlation between the number of times the Rehearsal tactic was used by a student and their final performance, this correlation is weak (r = 0.19, p = 8.82e-29), and much weaker compared to the other learning tactics.

### 3.4 Learning Strategies

Three groups of learners, known as learning strategies [24], have been identified based on the frequency of using the learning tactics discussed in Section 3.3. The identified learning strategies can be mapped to well-recognised learning approaches introduced by [37, 40, 41]. Therefore, we used these learning theories to name the detected learning strategies and describe them according to the available educational information.

The results show that the majority of students (73% of all learners) are **low engagement/surface** learners who only used two learning tactics, Elaboration and Problem-solving, during their interaction with the course. They achieved a low final grade (m=31, std = 27) and had low levels of engagement compared to the other two groups of learners (**Fig. 5**).

**Fig. 5.**
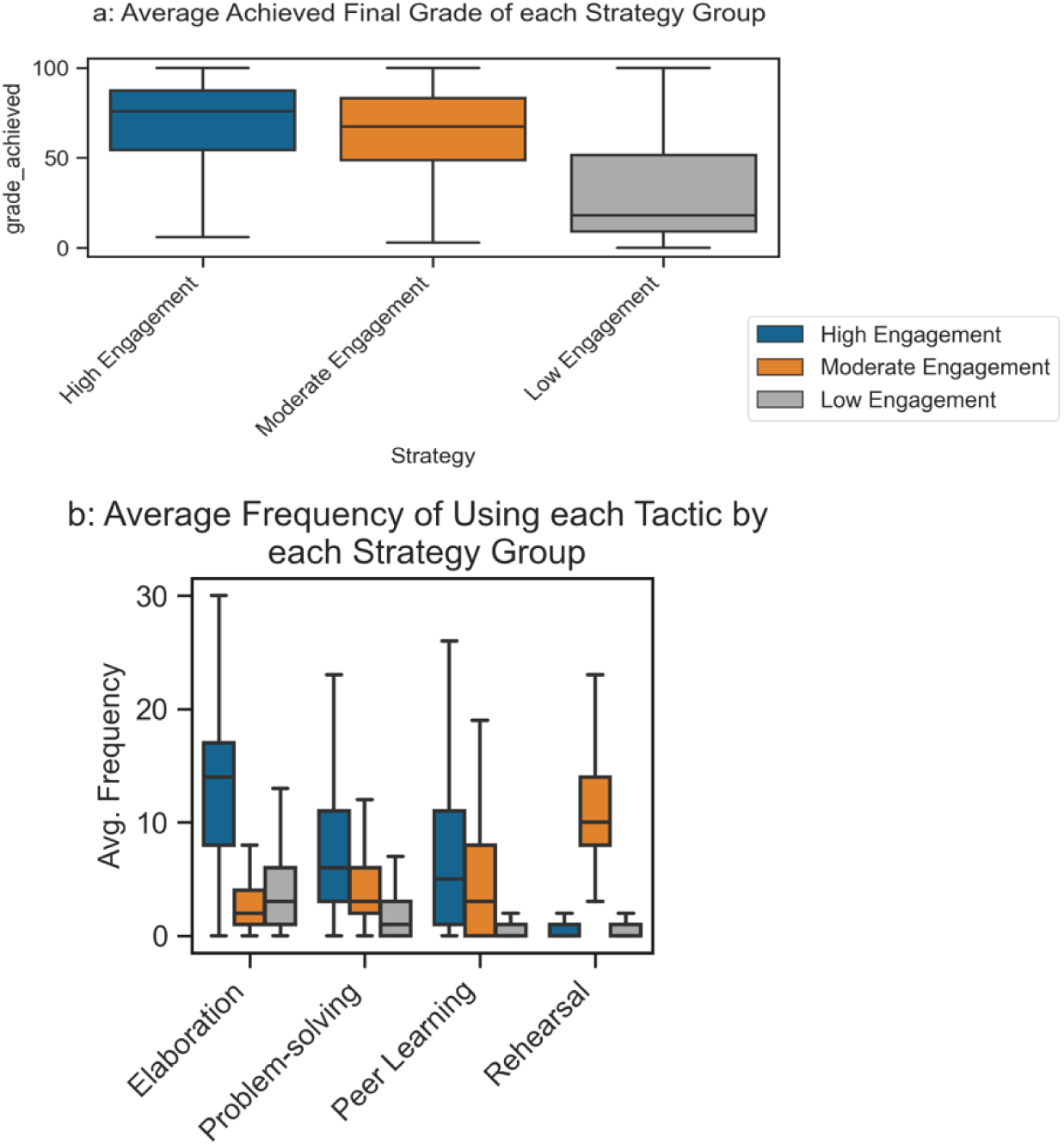
Average frequency of using each learning tactic by low, moderate, and high engagement learners (5b). as well as averaged final grade for each strategy group (5a).

The second group of students (19% of all learners), **high engagement/deep** learners, used all learning tactics except Rehearsal with higher frequency, as shown in **Error! Reference source not found.**This group has the highest frequency of using the Elaboration, Problem-solving, and Peer learning tactics. This group also achieved the highest final grade (m=68, std=25), whereas surface learners achieved the lowest grade and were overall not successful in passing the course.

The third learning strategy group (8% of all learners) are **moderate engagement/strategic** learners who employed all learning tactics with moderate frequency, except for Rehearsal, which was used with relatively higher frequency. It can be inferred that these students strategised their learning by regulating their time in such a way as to only revisit and seek important parts of video lectures in order to achieve an acceptable grade. Although moderate engagement learners used all four discovered tactics, their final performance (m = 62, std = 25) is lower than deep learners. Deep learners did not use the rehearsal learning tactic, but it appears that the elaboration tactic, along with the other two tactics, was enough for them to achieve higher grades than moderate-engagement students.

The Kruskal-Wallis test shows a significant association between the discovered learning strategies and student final performance (p = 2.51e-169, statistic = 776.43).

## 4 Discussion

This study employed artificial intelligence methods to provide insights into health data science education by analysing an MOOC with over 3,000 enrolled learners. The findings reveal that there is not a strong difference in the frequency of visits to reading materials, video lectures, and labs, although students tended to visit labs and lectures slightly more than reading materials. Also, based on the results, students who actively engaged with practical resources, such as labs, discussions, and projects, achieved higher final grades.

Furthermore, the results indicate that the students engaged more with the Sequence Processing and Python Programming topics. However, students moved forwards and backwards more in the programming topic videos compared to other topics. One inference is that students might find this topic challenging.

To analyse the students’ learning strategies, four learning tactics were identified that are in line with educational learning theories [38, 41]. The most frequently used tactic is *Elaboration*, which involves learning by pausing and replaying video lectures, possibly to contemplate the taught concepts or take notes. Based on the MSLQ learning theory [38], this tactic assists learners in retaining information in their long-term memory by establishing connections between the items that need to be memorised. This tactic includes pausing and replaying a video lecture, potentially in order to rephrase or condense information into a summary, make comparisons and take notes in an active manner. This tactic supports a learner in combining and linking new information to their existing knowledge.

The second most used tactic is *Programming and Problem-Solving*, where students engaged with the programming labs and solved the programming assignment. Given the interdisciplinary nature of health data science, students need to develop their knowledge in programming and improve their problem-solving skills [42]. Therefore, this tactic can be effective for students to apply their theoretically acquired knowledge to solve problems by using programming.

The third tactic is *Peer Learning,* which involves communicating with peers in the discussion forums and solving the peer-reviewed assignment. This tactic was found to have a positive correlation with students’ final performance, which was stronger than for the other learning tactics. Existing literature confirms that peer learning is associated with high performance, especially in online courses where students do not have the opportunity to discuss the learning materials face to face [38, 39]. It could be even more useful in the health data science field because students have diverse backgrounds; therefore, they can share their ideas and perspectives towards multi-disciplinary topics to develop in-depth knowledge and reflect on different approaches.

The final tactic is *Rehearsal*, which involves learning mostly by going forwards and backwards in video lectures instead of watching them from beginning to end. According to the MSLQ learning theory, it is a basic tactic for learning, which involves repeating information, again and again, to memorise it instead of deeply thinking about it. This tactic is effective for simple tasks and for calling information stored in the working memory, but not for acquiring new information that will be stored in the long-term memory. This tactic is believed to impact attention and the process of encoding information, but it does not seem to help students develop relationships between the information or integrate the information with their prior knowledge [38, 39]. Based on a recent systematic review [39], one study [43] showed that this tactic has a weak positive impact on student performance; while two studies [44, 45] did not find any significant association between rehearsal and performance. In this study, we found a weak correlation between rehearsal and student final grade (the weakest correlation compared to the other learning tactics). Interestingly, the Rehearsal tactic was not used much by deep learners, who achieved a higher final grade than surface and strategic learners.

Based on the frequency of using the learning tactics, three learning strategy groups were identified: low engagement/surface, high/engagement/deep, and moderate engagement/strategic. The learning strategies detected are highly accordant with the well-recognised learning approaches introduced by Biggs [41], Marton and Säljö [37], and Entwistle [40]. These scholars have described three learning approaches named deep, strategic, and surface learning, which are not the intrinsic characteristics of students [41], rather they are selected by students based on the task type and cognitive conditions. Also, students’ motivations and intuitions, the learning environment, the way the course is delivered, and the learning contents are the key factors that influence the choice of a learning approach by students [24, 40].

The high engagement/deep learners’ group is characterised by a high level of engagement, a high frequency of employing various tactics, and a high number of quizzes and project submissions, which is consistent with a deep learning approach, by which students engage with high frequency with the course materials, they are highly engrossed in the ideas and actively try to relate them to previous knowledge [38]. Previous studies have shown that adopting the deep learning approach results in better academic performance [24, 46]. Furthermore, the students with deep learning strategy obtained the highest marks in course assessments compared to other students, which indicates their in-depth knowledge. Also, the high use of the Elaboration, Problem-Solving, and Peer Learning tactics by these students reveals that they tend to focus on course materials for a long time (these tactics include long sessions), relate learning materials to their prior knowledge, focus on programming labs to solve problems and learn from peers and solve a project, which are all aligned with the characteristics of the deep learning approach.

The surface learning approach is adopted by students whose intention is to not fail and who want to achieve a passing mark rather than gain a deep understanding of the materials or obtain high marks. Therefore, these students mainly memorise the required information that is necessary for the exams, do not focus on abstract ideas, and mostly rely on details [37]. This approach has similar characteristics to the low engagement strategy in our study because the students using this strategy only used the Elaboration and Problem-Solving learning tactics with low engagement, resulting in low performance. They also did not use the Peer Learning tactic, which had the highest impact on student performance, because the peer-reviewed assignment corresponds to 30% of the final grade. In the DSM course, the passing score is 50 out of 100, and 50% of the final grade is related to quizzes. A deeper level of knowledge is required for the project compared to the quizzes. DSM students employing a surface approach tend to concentrate primarily on quizzes (by employing the problem-solving tactic) in order to achieve a passing score without investing significant effort in the project.

The strategic or achieving learning approach is described in educational theory as a combination of the surface and the deep approaches [46]. The main motivation of students adopting this approach is to get high scores and manage their efforts to make the most of the assessments done [47]. Therefore, they try to find the demands of assessments, manage their time, study in an organised manner, and routinely make sure that they use proper materials [47]. This learning approach is similar to the moderate engagement strategy in our study. The students with this strategy had moderate efforts, moderate frequency of using different tactics, and moderate performance in comparison to the two other strategies. They also mostly used the Rehearsal tactic, which shows that students moved forwards and backwards in video lectures instead of watching them from the start until the end. This is consistent with the characteristics of strategic learners who prefer to apply timely efficient tactics to manage their learning. Therefore, they used the Rehearsal tactic more than Elaboration because the Elaboration tactic is attributed to more effort, such as pausing and replaying videos instead of only seeking videos. This is also supported by the finding that the number of learning actions per learning session was higher in the Elaboration tactic.

It is worth pointing out that students use different learning strategies in different courses [48]. The learning tactics and strategies in health data science courses may differ from those in traditional biology or data science courses due to their interdisciplinary nature. Students in health data science must engage with both domain-specific biomedicine knowledge and data science concepts, requiring distinct strategies to facilitate their learning process [9, 49]. The learning tactics and strategies identified in this study for the health data science course are unique, though they do share some similarities with the tactics and strategies reported in previous studies on biology and computer programming courses [24].

### 4.1 Recommendations for Course Design and Education Improvement

The identified insights about health data science students can help to design better courses and programmes in this field. Most educational design models [50–52] need information about students to design effective pedagogical frameworks (e.g., pedagogical strategy and tactics) and educational settings (e.g., learning tasks and organisational forms). For example, learning tactics and strategies could be defined in the form of pattern languages based on [50] for designing better educational frameworks. In other words, a key implication of our study is to provide health data science educational designers with insights about HDS students and their learning behaviours that can potentially assist them in designing better educational courses and frameworks. Our recommendations based on this study are as follows:

In the DSM course, there is a peer-reviewed project in the last week that is responsible for 30% of the final grade. Since many students were not successful in submitting the final assignment, our recommendation is to invite students to work on the assignment throughout an HDS course rather than only in the last week. This can be particularly helpful for LP students, as it can encourage them to remain engaged during all weeks [53].

We showed that students in DSM engaged with a diverse range of learning resources (lab, reading, video, quiz, and project). Previous research has shown that utilising diverse learning resources, such as reading materials, interactive video lectures, games, labs, and so on, can enhance students’ learning experiences [54]. As an example, some students may prefer to look at reading materials instead of videos, or vice versa. Therefore, the available resources should be diverse, as students are diverse in HDS courses. Additionally, previous research shows that integrating interactive resources, such as gamification tools, may increase student engagement and lead to improving their learning outcomes [55, 56].

Our findings demonstrate that student engagement with topics decreased over the course, as evidenced by higher engagement with starting topics compared to ending topics. In the DSM course, as in many MOOCs [57], students have access to all topics/weeks upon enrolling on the course, which might overwhelm students given the large volume of learning materials. This can decrease their motivation, especially if they browse materials and assessments in the final topics and find them challenging. To address this issue, a potential solution could be to provide access to course material sequentially in such a way that a student can only have access to the subsequent topics upon the successful completion of previous topics.

Our results show that students had higher interaction with video lectures in the introductory Python programming topic compared to the other course topics, in particular higher video seek, pause, and play action. There are two possible explanations here. On one hand, students proficient in programming might have found the initial topic relatively straightforward and thus, did not engage with the entire video lecture from beginning to end. On the other hand, students with no programming background might have found the topic challenging and therefore rewatched certain parts of the videos. Given this mismatch, one might wonder how to best design a health data science course that works for diverse student backgrounds, including both computational and non-computational backgrounds. Our recommendation is to still provide introductory programming topics, but make them compulsory for students with no programming experience (so as to get up to speed with programming concepts) and optional for students with advanced programming skills (so that they are not disengaged). Once this is established as a baseline, subsequent programming-related tasks in the course should be designed at a balanced level of programming difficulty [58, 59].

Based on the findings, peer learning in HDS can help students to achieve higher performance. Therefore, grouping students in such a way that each group contains students with different backgrounds and asking them to work on a project may help them not only better learn both computational and medical aspects of the course, but also help them to learn how to collaborate in an interdisciplinary community, which is essential for a career in health data science [49, 60].

### 4.2 Recommendations for Teachers and Learners

The results of this study have implications not only for educational design, but also for learners and instructors. Learners sometimes are not aware of the most effective learning strategies, and informing them can possibly improve their future learning experiences [61–63]. However, course design is not the whole story, and teachers’ presentation approach also plays an important role in improving students’ learning outcomes. Therefore, we also provide some recommendations for teachers that may help them teach HDS more effectively.

#### Recommendations for Learners

Applying multiple learning tactics when interacting with a course was found to be more effective than only using one or two learning tactics. Our findings, similar to previous research [48]. For example, in order to achieve a good grade in programming and enhance one’s programming skills, simply watching video lectures about programming is not enough. Students who practised coding and used the discussion forums to ask questions and solve the peer-reviewed project were more successful.

The results indicate that successful students not only relied on required knowledge for assessments but also went beyond the syllabus [37] and even engaged with optional sections of the DSM course (e.g. case studies). Therefore, we recommend to students to not only follow the essential parts of a health data science course, but also study additional resources to get a comprehensive knowledge of each topic.

Our findings demonstrate that students who paused and replayed video lectures in order to relate the taught concepts to their prior knowledge, take notes, or think deeply about the topics were more likely to achieve high performance in DSM. Our recommendation to health data science students is, therefore, to use the Elaboration tactic along with other effective learning tactics (Peer Learning and Problem-Solving). Using the Rehearsal learning tactic without deep comprehension is not always effective.

#### Recommendations for Teachers

Previous studies [23, 64] have shown that personalised feedback can help students to improve their learning. We recommend that instructors consider students’ learning tactics, strategies, and preferences when they are providing feedback to them.

Our results also show that although there is a relatively low number of posts in the DSM discussion forums, many students visited the discussion forums to read other students’ questions and answers. Given our finding that students who engaged in discussions more were more successful, teachers should encourage students to participate in the discussion forums. Students might be introverted or feel uncomfortable posting on discussion forums; therefore, teachers should motivate them through the use of appropriate techniques. As an example, a study showed that the active presence of teachers in discussions, through asking questions and following up with additional questions, can enhance students’ engagement [65]. Therefore, specifically for HDS courses, we recommend that teachers post a question in the discussion forums and ask students to share their opinions. Also, posting about cross-disciplinary research findings related to each topic might encourage students because it can show the application of each topic [66].

### 4.3 Limitations and Directions for Future Work

Given that in this study we analysed one health data science course, further research is needed to validate the generalisability of our findings. Also, given that the DSM course is a self-paced MOOC, our findings might not apply to other online courses or face-to-face classes. This is particularly important when considering the fact that students who enrol on MOOCs have different motivations [67] and it is possible that some of them did not focus on assessments because it is not part of their mandatory study programme. This can impact findings related to student performance. We invite researchers to analyse the learning strategies employed by health data science students in other online or face-to-face courses. Unfortunately, we did not have access to temporal data (time spent to study each resource) for readings, discussions, and labs in the DSM course. Therefore, the student engagement with different topics (RQ2) was only explored based on the video lectures’ temporal data.

Our findings are limited to students’ click-stream data about the course on the Coursera platform. Since there are well-recognised survey tools, such as MSLQ [38] and self-regulation learning [68], for identifying students’ learning preferences and strategies that can uncover students’ perceptions about their learning regardless of the learning environment, it is worth collecting self-reported data and combining it with data-driven information as has been done for non-HDS courses [69], so as to strengthen results. We regard this as a fruitful avenue for future research.

## 5 Conclusions

Given that little is known about the learning behaviours and experiences of health data science students, conducting research to provide insight into health data science education is necessary. To address this important research gap, we employed artificial intelligence methods to analyse a health data science MOOC in order to understand students’ learning tactics, strategies, and engagement with learning materials and topics. We also provided suggestions supported by our findings for teachers, learners, and course designers in order to improve health data science education. The key findings of this study are the following:

- Students who engaged more with practical resources, such as projects, labs, and discussions achieved higher final grades.
- Among the topics taught, it seems that students were more engaged with Python Programming and Sequence Processing topics.
- The Elaboration tactic (connecting new information to their prior knowledge) was used more, and this tactic is effective for achieving high performance.
- The Peer Learning tactic had the highest correlation with the final grade.
- The Rehearsal tactic (memorising information by repeating) had the lowest correlation with the final performance, and deep learners, who are the most successful students, did not use this learning tactic.
- Students who employed a deep learning strategy utilised a range of different learning tactics throughout the course and engaged with all educational resources that enabled them in achieving higher final grades.

## List of Abbreviations

AHC: Agglomerative Hierarchical Clustering
AI: Artificial Intelligence
AVG: Average
DSM: Data Science in Stratified Healthcare and Precision Medicine
HDS: Health Data Science
HP: High Performance
LP: Low Performance
MOOCs: Massive Open Online Courses
MP: Moderate Performance
MSLQ: Motivated Strategies for Learning Questionnaire
NO: Number
RQ: Research Question

## Declarations

### Ethical approval and consent to participate

This study does not involve experiments with human participants. Therefore, informed consent to participate does not apply. We have used secondary data that includes learner (i.e. human) data. This data is anonymised and it was collected by the Coursera platform and shared with us in the University of Edinburgh for research purposes. We have received ethics approval for this research (which involves the use of this secondary data) by the Informatics Research Ethics Committee at the University of Edinburgh [application number: #88883]. Coursera collects this data in accordance with its Terms of Use and its Privacy Notice. According to Coursera’s Privacy Notice, by using the Coursera website, learners agree to the use of their data for research purposes. Coursera has shared this data with us following its Research and Data Sharing Policies, which protect learners’ right to privacy.

### Consent for publication

Not applicable.

### Availability of data and materials

The datasets generated and analysed during the current study are not publicly available due to ethical and legal restrictions. Data are however available from the Ethics Committee of the University of Edinburgh and the Coursera platform upon reasonable request.

### Authors’ contributions

All authors contributed to conceptualisation and design of the study. The implementation of the method as well as analysis of the results have been done by NR and supervised by AM, KG, and MG. The first draft of the paper was written by NR and improved by AM, KG, and MG. All authors read and accepted the final version of the paper.

## Acknowledgments

We would like to thank the Precision Medicine programme of the University of Edinburgh, as well as the Medical Research Council, for their support of this project aimed at enhancing health data science education. Additionally, we would like to express our appreciation to the Coursera platform and the students who participated in the course, whose contribution was invaluable to this research.

