## Supplementary file for "Providing Insights into Health Data Science Education through Artificial Intelligence"

### Method

The input learning sessions that had at least one of the following actions were considered for the analysis:

- **Video-start:** Starting to watch a video for the first time.
- **Video-play:** Playing a video lecture.
- **Video-end:** Watching a video until the end.
- **Video-seek:** Skipping forward or backwards throughout a video.
- **Video-pause:** Pausing a video lecture.
- **Video-revisit:** Watching a video for the second time or more.
- **Lab engagement**: Any activity related to programming and Jupyter notebook activities.
- **Reading engagement:** Any activity related to the reading material such as visiting reading pages.
- **Discussion engagement:** Any activity related to the discussion forums.
- **Quiz-visit:** Visiting quiz-related pages without submitting answers.
- **Quiz-failed:** Failing a quiz (score lower than 50% of total score).
- **Quiz-passed:** Passing a quiz.
- **Peer-reviewed project engagement:** Any activity related to the peer-reviewed projects, such as reviewing submissions of their classmates.

Learning sessions were profiled for each student and analysed to identify the students' learning tactics. To have a more representative dataset, multiple pre-processing steps (as used in the previous work [1]) such as removing outlier sessions were applied. Following the approach in previous work [1], learning tactics of the students were detected with the use of process mining and clustering methods. In particular, First-Order Markov Models method, as implemented in the pMineR package

[2], was employed to calculate the transition probability matrix of actions. The number of possible learning tactics (no. tactics=4) was estimated based on automatic algorithms such as Elbow method and Hierarchical clustering dendrogram as well as considering contextual knowledge. To identify the learning tactics, the Expectation-Maximisation algorithm [2] was applied to the obtained transition probability matrix.

A student may apply a range of tactics throughout a course. Therefore, a learning strategy is defined as the goal-driven usage of a collection of learning tactics with the aim of obtaining knowledge or learning a new skill [1]. To extract the various strategies adopted by students and following methods established in related work, the frequencies of using each tactic by each student were calculated and transformed to the standard normal distribution. Finally, Sessions with at least one coded action were selected. First-order Markov Model was applied to create a process map and a transition matrix for all pairs of actions. The transition matrix was used to cluster the sessions. The frequencies of detected learning tactics were used to cluster students into learning strategy groups the strategies (no. strategies=3) were identified by clustering the students using Agglomerative hierarchical clustering with Ward's linkage and Euclidean distance of the normalised vectors as the distance of students. To shed light on the identified learning tactics, the TraMineR package [3] was used for analysing the distribution, duration and order of employing learning actions.

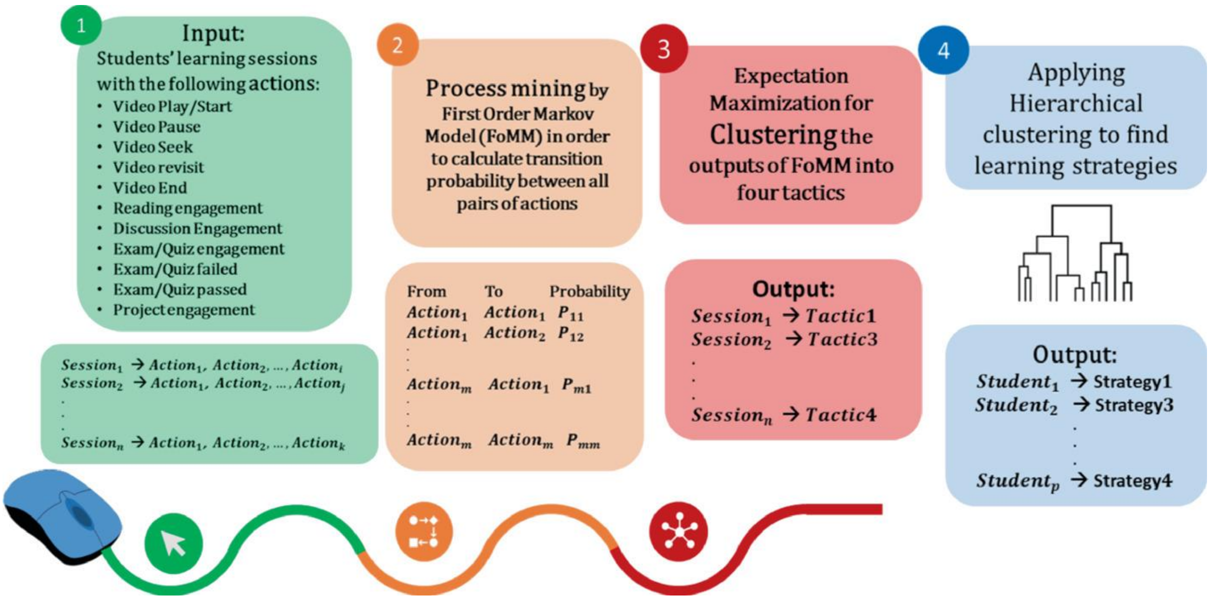


*Figure 1. Schema of the method: 1) Sessions with at least one coded action were selected. 2) First-order Markov Model was applied to create a process map and a transition matrix for all pairs of actions. 3) The transition matrix was used to cluster the sessions into four tactics using Expectation-Maximization method. 4) Hierarchical clustering was used to cluster students into four groups of strategies based on the frequency of their tactics. This figure was created by the first author and previously published at [4].*
